# Versatile roles of protein flavinylation in bacterial extracyotosolic electron transfer

**DOI:** 10.1101/2024.03.13.584918

**Authors:** Shuo Huang, Raphaël Méheust, Blanca Barquera, Samuel H. Light

## Abstract

Bacteria perform diverse redox chemistries in the periplasm, cell wall, and extracellular space. Electron transfer for these extracytosolic activities is frequently mediated by proteins with covalently bound flavins, which are attached through post-translational flavinylation by the enzyme ApbE. Despite the significance of protein flavinylation to bacterial physiology, the basis and function of this modification remains unresolved. Here we apply genomic context analyses, computational structural biology, and biochemical studies to address the role of ApbE flavinylation throughout bacterial life. We find that ApbE flavinylation sites exhibit substantial structural heterogeneity. We identify two novel classes of flavinylation substrates that are related to characterized proteins with non-covalently bound flavins, providing evidence that protein flavinylation can evolve from a non-covalent flavoprotein precursor. We further find a group of structurally related flavinylation-associated cytochromes, including those with the domain of unknown function DUF4405, that presumably mediate electron transfer in the cytoplasmic membrane. DUF4405 homologs are widespread in bacteria and related to ferrosome iron storage organelle proteins that may facilitate iron redox cycling within ferrosomes. These studies reveal a complex basis for flavinylated electron transfer and highlight the discovery power of coupling comparative genomic analyses with high-quality structural models.

## INTRODUCTION

Essential aspects of prokaryotic physiology take place beyond the bounds of the cell cytosol. Extracytosolic interactions can occur at the extracytosolic side of the inner membrane, periplasm, cell wall, or surrounding environment. Within the extracytosolic environment, redox reactions (defined by the reduction of an electron acceptor and oxidation of an electron donor) represent an important class of activities that have functions in respiration, maintenance/repair of extracytosolic proteins, and assimilation of minerals (Schröder, Johnson, and de Vries 2003; Bertini, Cavallaro, and Rosato 2006; Cho and Collet 2013).

Flavins are a group of small molecules that contain a conserved redox-active isoalloxazine ring system. Most microbes synthesize riboflavin (or vitamin B2), flavin mononucleotide (FMN) and flavin adenine dinucleotide (FAD). FMN and FAD serve as common cofactors within diverse redox-active enzymes (Fraaije and Mattevi 2000). Flavinylation describes the covalent attachment of a flavin moiety to a protein and frequently occurs in proteins involved in extracytosolic electron transfer (Bogachev, Baykov, and Bertsova 2018). The alternative pyrimidine biosynthesis protein, ApbE, is a widespread FMN transferase that flavinylates a conserved [S/T]GA[**S/T**]-like sequence motif (flavinylated amino acid in bold) within substrate proteins (Bertsova et al. 2013). ApbE-flavinylated proteins are integral for a number of extracytosolic electron transfer systems. These include the cation-pumping NADH:quinone oxidoreductase (Nqr) and Rhodobacter nitrogen fixation (Rnf) complexes, nitrous oxide and organohalide respiratory complexes, and a Gram-positive extracellular electron transfer system (Backiel et al. 2008; Buttet et al. 2018; Light et al. 2018; Zhang et al. 2017; Zhou et al. 1999). ApbE-flavinylation has also been implicated in electron transfer for a large family of flavin reductases that use different electron acceptors to power anaerobic respiration (Bogachev et al. 2012; Kees et al. 2019; Light et al. 2019; Little et al. 2023).

Using the presence of ApbE and/or FMN-binding domains as genomic markers, we previously screened 31,910 genomes representative of the diversity of prokaryotic life and found that ∼50% encoded machineries involved in flavinylation (Méheust et al. 2021). We observed that ∼50% of genomes with ApbE flavinylation machineries lacked one of the previously characterized systems mentioned above. By mining the gene colocalization dataset, we discovered that extracytosolic flavinylation occurs in proteins with a variety of different domain topologies and is associated with novel transmembrane components that link redox pools in the membrane to the extracytosolic space (Méheust et al. 2021). Previous studies thus provided clear evidence that ApbE flavinylation is a central component of various prokaryotic extracytosolic redox activities but little insight into the molecular basis of its function.

The recent development of AlphaFold, an artificial intelligence-powered protein structure prediction tool, has created new opportunities for high-throughput analysis of protein structures with great speed and accuracy (Jumper et al. 2021; Evans et al. 2021). In this study, we used computational modeling and comparative genomic approaches to comprehensively evaluate the predicted functions and structures of flavinylated proteins. Our results reveal a high degree of diversity in structures of flavinylated proteins, highlighting the versatile roles that flavinylation may play in bacterial biology.

## RESULTS

### AlphaFold models reveal the diverse context of ApbE flavinylation

To address general principles of electron transfer through ApbE-flavinylated proteins, we generated representative AlphaFold models of previously identified ApbE flavinylated domains (FMN-bind, NqrB/RnfD, and DUF2271) and compared these to experimentally characterized structures. The resulting collection of structures reveals a diverse context of flavinylation that varies across the different classes of flavinylated proteins. Flavinylated FMN-bind, NqrB/RnfD, and DUF2271 domains each possess a distinct fold and a unique structural context of the flavinylation site (**Figure 1**). FMN-bind and DUF2271 are small soluble and generally non-descript domains with distinct folds (**Figure 1A & D**). NqrB/RnfD are transmembrane proteins which typically serve as subunits within larger multi-protein complexes in the cytosolic membrane (**Figure 1C**) (Hayashi et al. 2001). Analyses of structural models of confirmed flavinylation substrates thus provide evidence of a variable context of flavinylation sites that partially reflects distinctions in domain cellular localization between the cytosolic membrane, the periplasm, and the outer membrane.

**Figure 1.**
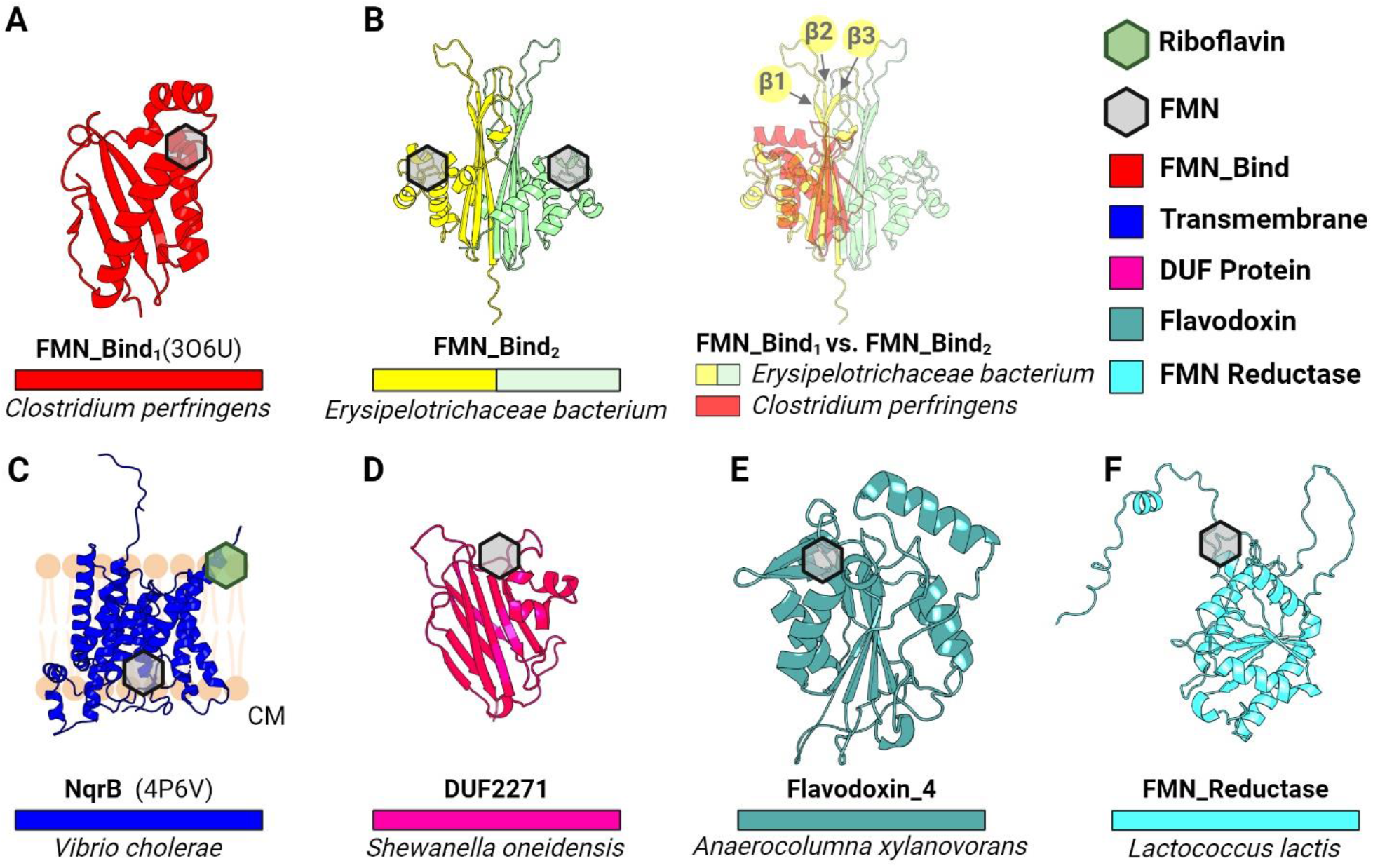
Structural context of ApbE flavinylation sites. (A) Previously resolved crystal structure of a flavinylated monomeric protein from *C. perfringens* (FMN_Bind_1_; PDB: 3O6U). (B) AlphaFold-predicted model of a double-flavinlated protein from *Erysipelotrichaceae bacterium* containing 2 FMN_bind_2_ domains (left) and its structural alignment with FMN_bind_1_ at the β-sheet face (right). Arrows highlight β1, β2, and β3 strands. (C) Previously resolved crystal structure of the B subunit from the Nqr complex (PDB: 4P6V). (D) AlphaFold-predicted mode of a novel AbpE substrate in *S. oneidensis*, namely DUF2271. (E-F) AlphaFold-predicted models of flavinylated Flavodoxin_4 (E) and flavinylated FMN reductase (F). Hexagons indicate predicted or confirmed locations of cofactors in structures. EC: extracytosolic space. PM: periplasm. CM: cytoplasm.

We also observed that structural distinctions distinguish domains from different proteins within ApbE substrate classes. For example, the FMN-bind domain, which is the most common flavinylated domain and is found in multiple characterized electron-transferring complexes (as NqrC, RnfG, and PplA), can be divided into two distinct structural subtypes (Borshchevskiy et al. 2015; Backiel et al. 2008; Light et al. 2018). The more common FMN-bind_1_subtype comprises a compact ∼120 amino acids structural core. The ∼160 amino acid FMN-bind_2_domain is less common and sometimes found in multiple copies within proteins. Proteins with multiple FMN-bind_2_ domains often contain an even number of domains (2 or 4) and an AlphaFold structure of the *Erysipelotrichaceae bacterium* 2 FMN-bind_2_ domain protein provides an explanation for this pattern. Relative to the FMN-bind_1_ domain, FMN-bind_2_ domains have insertions between β1-/β2-strands and the β3-strand/α1-helix that extends the β-sheet face of the domain (**Figure 1B**). This β-sheet face is predicted to interact with a homologous β-sheet on a neighboring FMN-bind_2_ domain within multi-FMN-bind_2_ domain proteins to produce a pseudo-symmetrical unit with two flavinylated sites (**Figure 1B**). Distinctions in FMN-bind_1_ and FMN-bind_2_ domain sequence thus seem to establish unique 1- and 2-flavinylated structural units, respectively. In turn, these structural differences may reflect distinct mechanisms of electron transfer.

### AlphaFold structures provide evidence of ApbE flavinylation evolving from non-covalent flavoproteins

To further expand our analyses of ApbE flavinylation substrates, we performed comparative genomic analyses to mine the Genome Taxonomy Database (GTDB) collection of 47,894 diverse bacterial and archaeal genomes and metagenome-assembled genomes (Parks et al. 2018). Through this analysis, we identified two groups of candidate flavinylation substrates that colocalize with *apbE* genes and contain an ApbE-like flavinylation motif sequence. Both candidates are related to characterized flavoproteins that contain a non-covalently bound flavin cofactor. The first group of proteins are part of the FMN_red protein family (Pfam accession PF03358) and have an extended C-terminal region that contains 1-2 flavinylation motif-like sequences (**Figure 1F & 2G**). The second group of proteins are part of the Flavodoxin_4 protein family (Pfam accession PF12682) and have an insertion with flavinylation motif-like sequence internal to the Flavodoxin_4 domain (**Figure 1E & 2A**). ApbE-associated proteins from both the Flavodoxin_4 and FMN_red protein families are predominately encoded by members of the Firmicutes phylum (**Figure 2E & 2K**).

**Figure 2.**
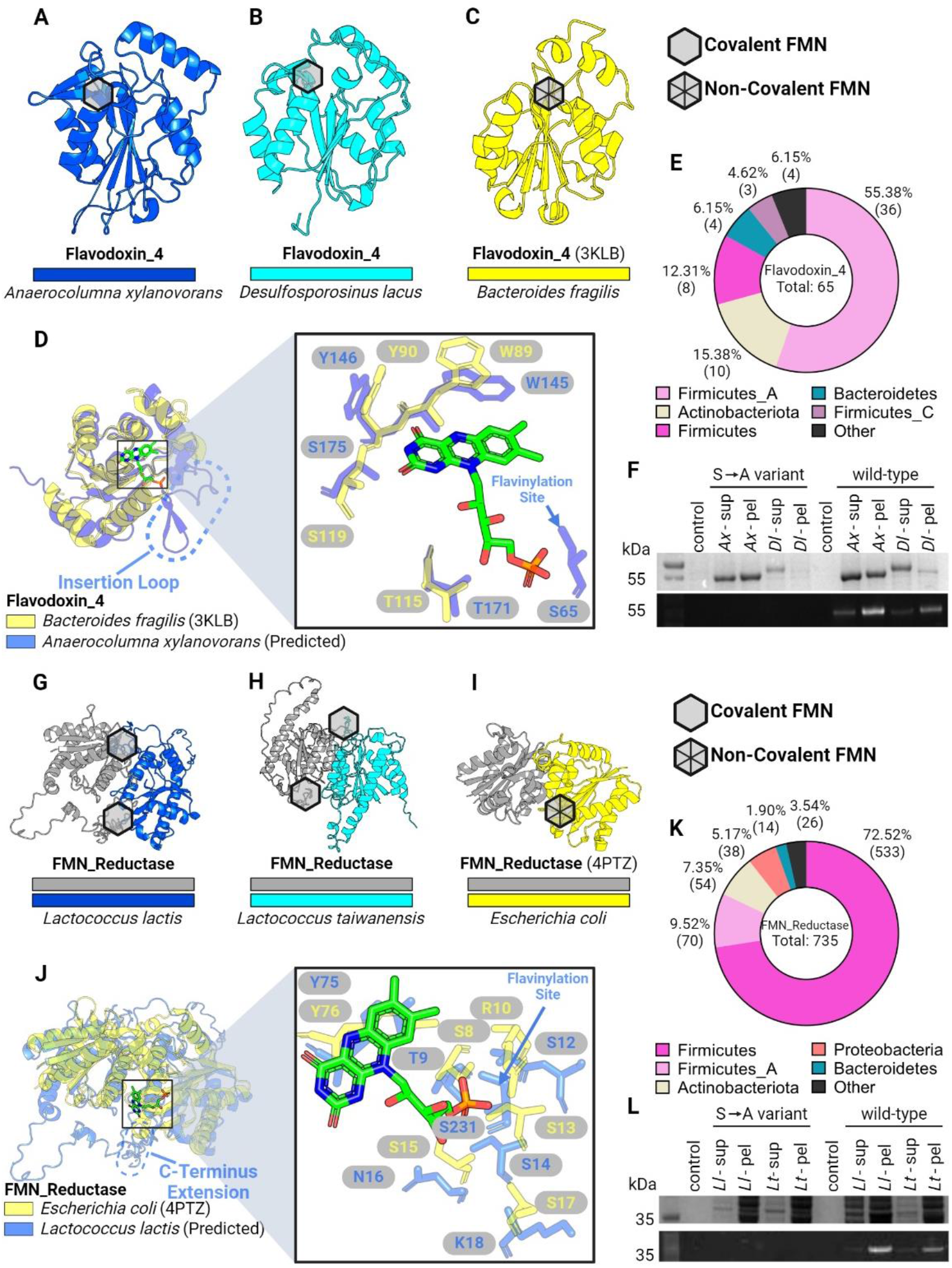
ApbE flavinylation evolved from non-covalent flavoproteins. (A-B) AlphaFold-predicted models for flavinylated Flavodoxin_4 proteins from *A. xylanovorans* (A) and *D. lacus* (B). (C) Previously resolved crystal structure of a Flavodoxin_4 with non-covalent FMN-binding (PDB: 3KLB). (D) Structural alignments of Flavodoxin_4 proteins with and without covalent FMN binding. Dashed loop highlights an inserted loop that is lacking in 3KLB (left). Right panel shows zoom-in view of non-covalent FMN molecule from 3KLB and surrounding residues. Arrow indicates the serine residue in flavinylated Flavodoxin_4 responsible for covalent FMN-binding. (E) Taxonomic distribution of Flavodoxin_4 proteins in the GTDB. (F) SDS-PAGE gel image of purified flavinylated Flavodoxin_4 from *A. xylanovorans* and *D. lacus*, visualized under UV. Covalent FMN moieties appear as bright bands. Flavinylation-deficient mutants were generated by converting FMN-binding Ser residues to Ala. (G-H) AlphaFold-predicted models for flavinylated FMN reductases in *L. lactis* and *L. taiwanensis*. (I) Previously resolved crystal structure of a FMN reductase with non-covalent FMN-binding (PDB: 4PTZ). (J) Structural alignments of FMN reductase proteins with and without covalent FMN binding. Dashed loop highlights a C-terminus extension that is lacking in 4PTZ (left). Right panel shows zoom-in view of non-covalent FMN molecule from 4PTZ and surrounding residues. Arrow indicates the serine residue in flavinylated FMN reductase responsible for covalent FMN-binding. (K) Taxonomic distribution of FMN reductases in the GTDB. (L) SDS-PAGE gel image of purified flavinylated Flavodoxin_4 proteins from *L. lactis* and *L. taiwanensis*, visualized under UV. Covalent FMN moieties appear as bright bands. Flavinylation-deficient mutants were generated by converting FMN-binding Ser residues to Ala. Hexagons indicate locations of FMN moieties in structure (empty: covalent; with diagonal lines: non-covalent).

To assess the likelihood of identified flavinylation motif-like sequences representing *bona fide* flavinylation sites, we compared ApbE-associated Flavodoxin_4 and FMN_red models to crystal structures of homologous proteins bound to a non-covalent flavin cofactor. For both Flavodoxin_4 and FMN_red structures, we observed that the core flavin-binding domain is structurally similar irrespective of putative flavinylation status (**Figure 2**). AlphaFold models of the ApbE-associated Flavodoxin_4 proteins from *Anaerocolumna xylanovorans* (**Figure 2A**; NCBI accession SHO45324.1) and *Desulfosporosinus lacus* (**Figure 2B;** NCBI accession WP_073032509.1) resemble a crystal structure of a homologous non-covalent flavin-binding protein (PDB: 3KLB) from *Bacteroides fragilis* (**Figure 2C**). However, the ApbE-associated Flavodoxin_4 proteins include an insertion between the 1β-strand and 1α-helix that contains the predicted flavinylated serine (**Figure 2D**). Strikingly, the predicted flavinylated serine/threonine in *A. xylanovorans* Flavodoxin_4 is perfectly positioned for the covalently bound flavin to engage the conserved flavin-binding site (**Figure 2D**).

A similar pattern is evident in the ApbE-associated FMN_red proteins. The crystal structure of an *Escherichia coli* FMN_red protein (PDB: 4PTZ) reveals a homodimer with symmetric flavin binding sites at the dimerization interface (**Figure 2I**). AlphaFold-multimer structures of the ApbE-associated FMN_red proteins from *Lactococcus lactis* (**Figure 2G;** NCBI accession WP_021723379.1) and *Lactococcus taiwanensis* (**Figure 2H;** NCBI accession WP_205872264.1) reveal a similar dimerization mode. Within the *L. lactis* model, predicted flavinylation sites are positioned on an unstructured C-terminal extension proximal to the conserved flavin-binding sites (**Figure 2J**). AlphaFold models thus provide evidence of the structural congruity between and ApbE-associated FMN_red and Flavodoxin_4 flavinylation motif-like sequences and structurally conserved flavin-binding sites.

As our analysis of AlphaFold structures suggested that flavinylation sites could secure flavins within established flavin-binding sites, we sought to address whether ApbE-associated FMN_red and Flavodoxin_4 domains were novel flavinylation substrates. We co-expressed *A. xylanovorans* and *D. lacus* Flavodoxin_4 proteins, as well as *L. lactis and L. taiwanensis* FMN_red proteins with their cognate *apbE* in *E. coli*. To address the specificity of flavinylation, we also expressed variants of these proteins with alanine point mutations at the predicted flavinylation site. SDS-PAGE analyses confirmed that FMN_red and Flavodoxin_4 proteins were flavinylated and that this required a serine/threonine at the predicted flavinylation site (**Figure 2F&L**). These findings thus highlight the utility of AlphaFold models in guiding protein function predictions, expand the repertoire of ApbE substrates, and suggest that, at least in some instances, flavinylated proteins evolved through the acquisition of a flavinylation motif at a preexisting flavin-binding site.

### Flavinylation-associated protein structures suggest diverse transmembrane electron transfer mechanisms

We previously identified five characterized and five uncharacterized electron transfer systems that co-localize on bacterial genomes with flavinylated proteins and which presumably utilize distinct mechanisms to mediate electron transfer from cytosolic or membrane donors to extracytosolic flavinylated domains (Méheust et al. 2021). Structures of the Nqr complex, the Rnf complex, and the *Pseudomonas aeruginosa* PepSY-like protein FoxB have been experimentally characterized, but the structural basis of other flavinylated-associated membrane electron transfer domains remains unknown (Steuber et al. 2014; Kishikawa et al. 2022; Josts et al. 2021; Vitt et al. 2022; Zhang and Einsle 2022). To clarify the context of transmembrane electron transfer, we generated AlphaFold or AlphaFold-multimer models for representatives of the remaining systems (**Figure 3A-3C & 4A-4C**). A comparison of the resulting models suggests a striking diversity in the structure and mechanism of flavinylation-associated transmembrane electron transfer which is further explored below.

**Figure 3.**
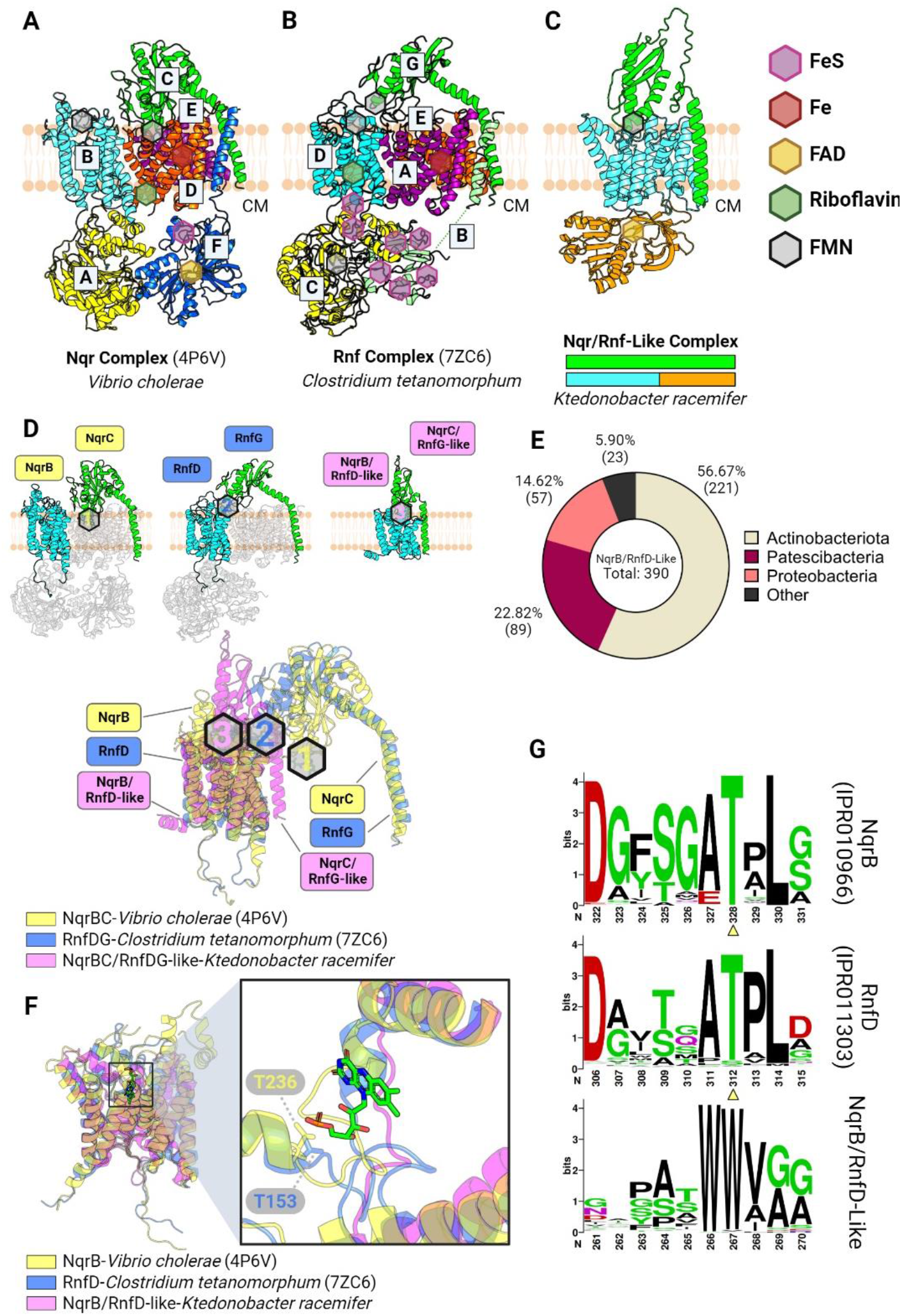
Flavinylation-associated transmembrane. (A-B) Previously resolved crystal structures of the Nqr complex (A; PDB: 4P6V) and the Rnf complex (B; PDB: 7ZC6). (C) AlphaFold-multimer model of the Nqr/Rnf-like complex containing a NqrB/RnfD-like transmembrane subunit and a NqrC/RnfG-like membrane-anchored extracytosolic subunit. (D) Structural alignment of Nqr, Rnf, and Rnf-like complexed based on the NqrB, RnfD, and NqrB/RnfD-like subunits. Subunits that do not share homology to NqrB/RnfD or NqrC/RnfG subunits are transparent (top). Predicted or confirmed FMN moieties are highlighted by hexagons, color-coded by their corresponding complex (bottom). (E) Taxonomic distribution of the NqrB/RnfD-like complexes in the GTDB. (F) Structural alignment of NqrB, RnfD, and NqrB/RnfD-like subunits (left) with zoom-in view showing the FMN moiety from NqrB, and the Thr residue in NqrB or RnfD responsible for FMN-binding (right). (G) HMM logos showing conserved Thr residues in NqrB and RnfD, but not in the NqrB/RnfD-like subunit. Hexagons indicate predicted or confirmed locations of cofactors in structures. CM: cytoplasm.

### Nqr, Rnf, and Nqr/Rnf-like complexes exhibit distinct but related paths for electron transfer

Previously reported structures of the six subunit Nqr complex reveal a semicircular electron transfer pathway in which electrons travel from cytosolic NADH to the extracytosolic NqrC flavinylation site, to the quinone terminal electron acceptor on the cytosolic side of the membrane (**Figure 3A**) (Steuber et al. 2014; Kishikawa et al. 2022). This reaction is coupled to the transfer of ions across the membrane and the creation of an electromotive force. Rnf is evolutionarily related to Nqr and possesses 4 homologous subunits but distinct substrate- and product-binding subunits that enable electron transfer between ferredoxin and NAD^+^. Recently reported cryoelectron microscopy structures provide evidence that RNF possesses a similar structure and mechanism as Nqr (**Figure 3B**) (Vitt et al. 2022; Zhang and Einsle 2022).

Our previous study identified Nqr/Rnf-like complexes as a distinct group of flavinylation-associated transmembrane subunits related to Nqr and Rnf (**Figure 3C & 3E**) (Méheust et al. 2021). Nqr/Rnf-like gene clusters contain only two apparent subunits. One Nqr/Rnf-like subunit is homologous to NqrC/RnfG subunits in Nqr/Rnf, respectively. The second subunit has an N-terminal membrane domain homologous to NqrB/RnfD from Nqr/Rnf and a cytosolic C-terminal NAD-binding domain (Pfam accession PF00175). In contrast to the semicircular Nqr and Rnf electron transfer path described above, we previously proposed that Nqr/Rnf-like complexes unidirectionally transfer electrons from NAD(P)H to extracytosolic electron acceptors (Méheust et al. 2021).

To gain insight into function of the Nqr/Rnf-like complex, we used AlphaFold-multimer to model the *Ktedonobacter racemifer* Nqr/Rnf-like complex. The high confidence AlphaFold-multimer model predicts that the subunit homologous to NqrC/RnfG and NqrB/RnfD intimately interact within the Nqr/Rnf-like complex. This interaction is noteworthy because corresponding subunits do not directly interact in the Nqr or Rnf complexes (**Figure 3A-3D**). While NqrC/RnfG and NqrB/RnfD subunits are both flavinylated in Nqr and Rnf, we observed that the flavinylation site in the structurally related NqrB/RnfD-like subunit is not conserved in the Nqr/Rnf-like complex (**Figure 3F&3G**). Strikingly, the unique interaction between NqrC/RnfG-like and NqrB/RnfD-like subunits in the Nqr/Rnf-like AlphaFold model brings the NqrC/RnfG-like subunit’s flavinylation site in close proximity of the apparent non-covalent flavin-binding site in the NqrB/RnfD-like subunit. This observation suggests that flavin covalently bound to the NqrC/RnfG-like subunit may conformationally sample the non-covalent flavin-binding site in the NqrB/RnfD-like subunit (**Figure 4D**). Collectively, these observations suggest a dynamic evolutionary history resulted in marked functional distinction between related Nqr, Rnf and Nqr/Rnf-like systems.

**Figure 4.**
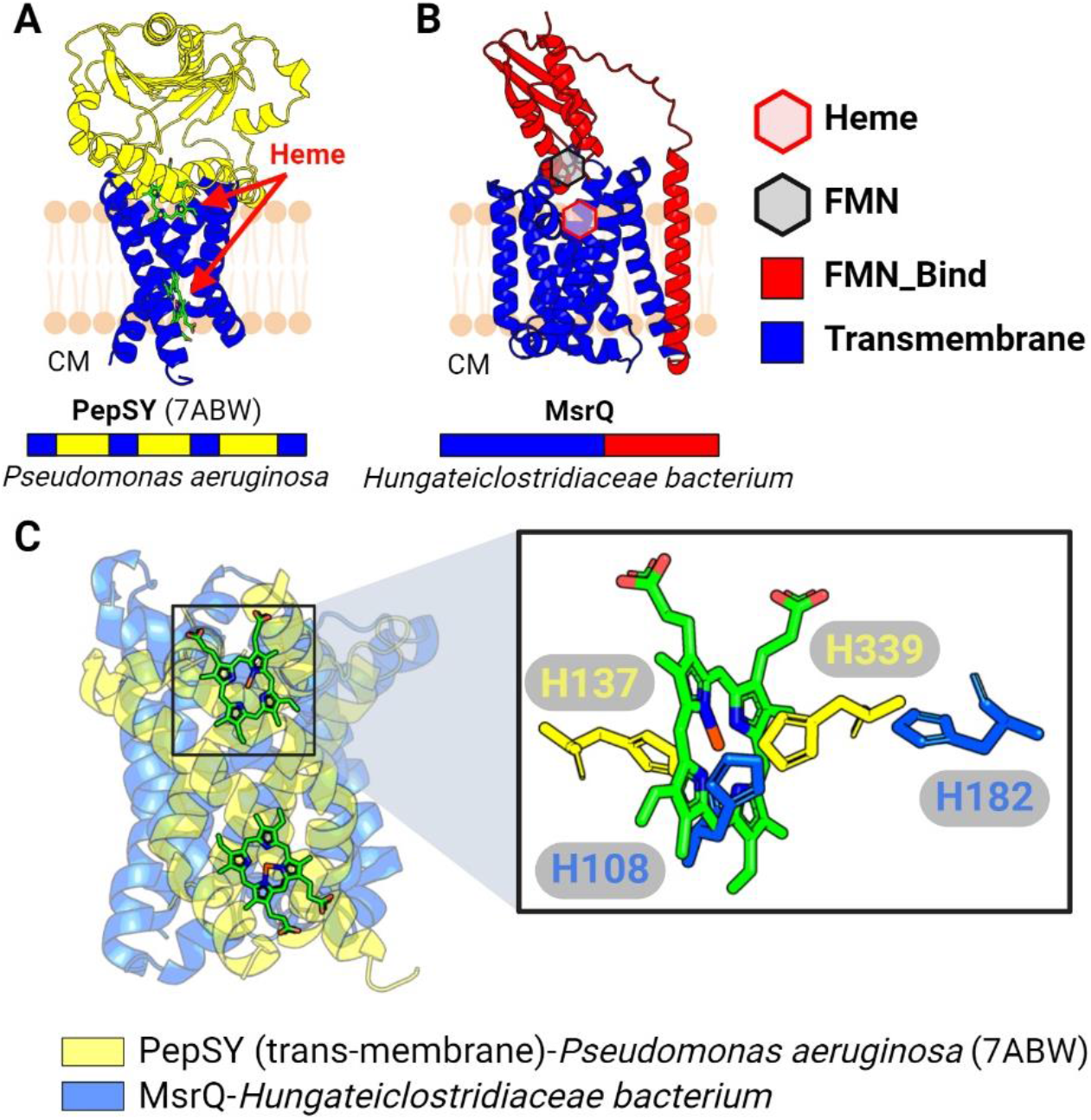
Membrane cytochrome complex associated with flavinylated proteins. (A) Previously resolved crystal structure of the PepSY complex (PDB: 7ABW). (B) AlphaFold-predicted structure of a complex containing the transmembrane cytochrome MsrQ and a membrane-anchored extracytosolic flavinylated protein. (C) Structural alignment between transmembrane segments of PepSY and MsrQ (left) with zoom-in view on the heme from PepSY and His residues from PepSY and MsrQ that are responsible for heme-binding. Hexagons indicate predicted or confirmed locations of cofactors. CM: cytoplasm.

### Structural similarities unite flavinylation-associated transmembrane cytochromes

PepSY-like and MsrQ-like transmembrane proteins are predicted to contain heme cofactors that transfer electrons across membranes. A recently reported crystal structure of the *Pseudomonas aeruginosa* PepSY-like protein revealed that it has two heme-binding sites and that each site contains two highly conserved histidines that coordinate heme binding (Josts et al. 2021) (**Figure 4A**). Despite low sequence identity, MsrQ-like AlphaFold structures exhibit considerable structural homology to the *Pseudomonas aeruginosa* PepSY-like protein, including two highly conserved histidines that come together to form a similar predicted heme-binding site (**Figure 4B & 4C**). These structures thus demonstrate a similar transmembrane core that is conserved within the flavinylation-associated cytochrome electron transfer apparatuses.

### DUF4405-like proteins establish a widespread class of flavinylation- and ferrosome-associated cytochromes

Having defined the structural features responsible for flavinylation-associated membrane electron transfer, we wondered if these insights could be leveraged to enable the discovery of novel proteins with analogous functionalities. We reasoned that such proteins would likely localize to genes clusters that contain *apbE* but lack a characterized flavinylation-associated electron transfer mechanism. We performed comparative genomic analyses mining *apbE* gene clusters that lack a known electron transfer mechanism and found that a transmembrane protein with a DUF4405 domain of unknown function frequently colocalized with *apbE* in Proteobacteria and Firmicutes species (**Figure 5B**). Consistent with these DUF4405s functioning in flavinylation-based electron transfer, we observed gene clusters containing *apbE* and DUF4405 genes also often encoded genes for a flavinylated Flavodoxin and a transporter that provisions *Listeria monocytogenes* with extracytosolic flavins (**Figure 5C**) (Rivera-Lugo et al. 2023).

**Figure 5.**
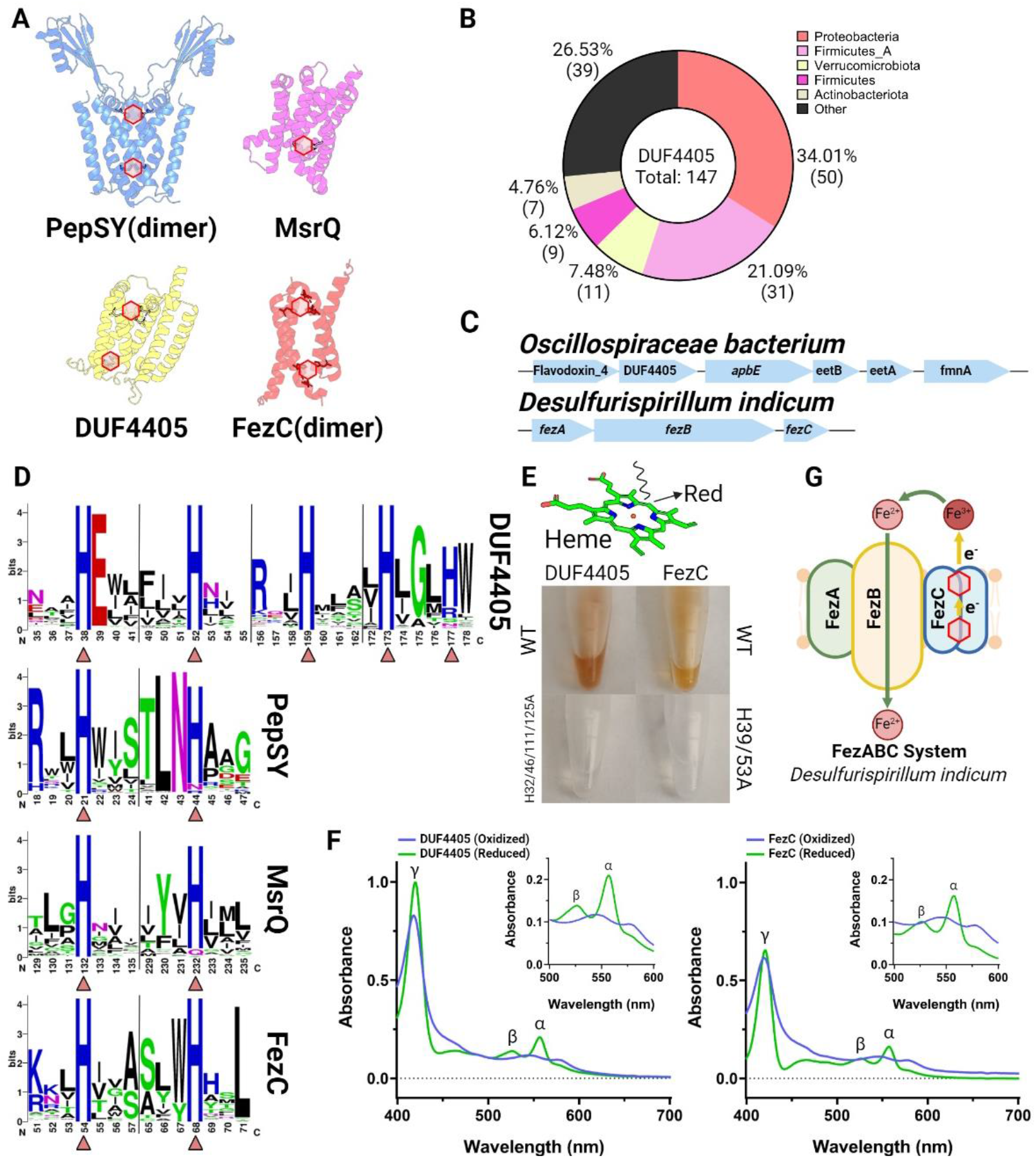
DUF4405 and FezC are novel cytochromes. (A) AlphaFold or AlphaFold-multimer models of MsrQ (homodimer), DUF4405, and FezC (homodimer). Histidine residues (His) responsible for heme binding are highlighted. (B) Taxonomic distribution of DUF4405 genes contained in a cluster with *apbE* in the GTDB. (C) Operons in *O. bacterium* or *D. indicum* encoding DUF4405 or FezC, respectively. (D) HMM logos showing conserved heme-binding His residues in DUF4405, PepSY, MsrQ, and FezC. Triangles highlight His residues shown in (A). (E) Visible red color from heme present in purified DUF4405 and FezC proteins. (F) UV spectrums of DUF4405 (top) and FezC (bottom) showing absorption peaks characteristic of heme B binding. (G) Possible role of FezC in iron transport within ferrosomes. Hexagons indicate predicted or confirmed locations of cofactors in structures.

To assess whether identified DUF4405 proteins might be involved in transmembrane electron transfer, we analyzed AlphaFold structures of representative proteins (**Figure 5D**). These structures revealed that DUF4405 possesses a six transmembrane structure with four centrally located histidines arranged strikingly similarly to PepSY-like and MsrQ-like structures (**Figure 5D & 5E**). Despite a low overall sequence homology to PepSY-like or MsrQ-like proteins, an inspection of sequence conservation within the DUF4405 domain revealed that its centrally located histidines are similarly the most highly conserved amino acids (**Figure 5E**). These analyses thus establish that DUF4405 resembles flavinylation-associated cytochromes in its pattern of conserved histidine placement.

To test the hypothesis that DUF4405 represents a novel class of cytochromes, we recombinantly expressed the *Oscillospiraceae bacterium* DUF4405 protein. Strikingly, DUF4405 overexpression conferred *E. coli* cells with a pinkish hue typical of heme-binding protein (**Figure 5F**). Purified DUF4405 retained this color and spectroscopic analyses revealed absorbance peaks consistent with heme B binding (**Figure 5H**). These results thus demonstrate that DUF4405s represent a novel class of cytochromes frequently associated with flavinylation-associated electron transfer.

### FezC is a ferrosome cytochrome

As only a minority of proteins with DUF4405 domains colocalize with *apbE*, we wondered whether the identification of DUF4405 as a cytochrome might clarify the function of other DUF4405 proteins. The most relevant-seeming mention of DUF4405 within the scientific literature noted that the *Desulfovibrio magneticus* ferrosome protein FezC possesses sequence homology to DUF4405 proteins (Grant et al. 2022). Ferrosomes are recently discovered membrane-enclosed organelles that act as an intracellular store of iron within some bacteria (Grant et al. 2022; Pi et al. 2023). As flavinylated systems commonly facilitate iron transport across membranes and this activity could be directly relevant for ferrosomes (which contain a putative ferrous iron transporter, FezB), we reasoned that the functionally uncharacterized FezC might be a DUF4405-like cytochrome. Indeed, an AlphaFold model revealed that FezC could form DUF4405-like heme-binding sites via homodimerization and recombinant FezC exhibited cytochrome-like properties similar to DUF4405 (**Figure 5F & 5H**). These results establish that FezC is a cytochrome and suggest that it may play a role in modulating iron redox status to facilitate iron transport into and/or out of ferrosomes (**Figure 5G**).

## DISCUSSION

The importance of AbpE flavinylation for prokaryotic extracytosolic redox activities has become increasingly apparent in recent years. In this study we combine comparative genomic context analysis of flavinylation-associated gene clusters with AlphaFold structural modeling to explore the molecular basis of flavinylation-associated electron transfer. Our findings showcase how recent advances in protein structural modeling enabled by AlphaFold can be leveraged for discovery and provide evidence that ApbE flavinylation is involved in a wide range of cellular processes.

By examining structural models of proteins encoded in flavinylation-associated gene clusters lacking a predicted transmembrane electron transfer apparatus, we identify DUF4405 as a putative electron-transferring cytochrome. Broadening these analyses, we find that related cytochromes include ferrosome components with obvious potential roles in modulating the redox state in these iron-containing organelles. These findings highlight how the iterative application of comparative genomic analyses and structural modeling can enable unpredictable protein functional attributions.

Our structural analysis of proteins encoded on flavinylation-associated gene clusters led to the discovery of two classes of ApbE flavinylated proteins (Flavodoxin and FMN_red) that are closely related to unflavinylated flavoproteins (i.e., which non-covalently bind their flavin cofactor). These findings have implications for our understanding of the evolution and significance of protein flavinylation, demonstrating that, at least in some cases, flavinylation may have emerged as an evolutionary addition to unflavinylated precursor proteins. Moreover, the observation that extracytosolic Flavodoxin proteins exhibit signs of flavinylation (in contrast to unflavinylated cytosolic members of the family) is consistent with the main role of ApbE flavinylation being to prevent flavin diffusion and loss in extracytosolic space.

In summary, our study demonstrates how comparative genomic analyses coupled with AlphaFold protein structure analyses can be leveraged to infer novel protein functions. Our results provide new insight into the structural context of ApbE flavinylation and suggest that this modification may play a broad role in bacterial biology. Future experiments will be needed to fully understand the function of ApbE flavinylation and its role in bacterial physiology.

## METHODS

### Identification of flavinylated protein candidates

The three flavinylated protein candidates FMN_red, DUF4405 and Flavodoxin were identified by searching their Pfam accession number (PF03358.18, PF14358.9 and PF12682.10 respectively) in the proteomes from 47,894 functionally annotated bacterial and archaeal genomes from the Genome Taxonomy Database (GTDB, release 202) (Parks et al. 2018). FMN-bind_2_ domains were through blast searches. Briefly, protein sequences were functionally annotated based on the Pfam accession number (Pfam database version 33.0) (Mistry et al. 2021) of their best match using Hmmsearch (E-value cutoff of 0.001, version 3.3.2) (Eddy 1998). The five genes downstream and upstream of genes FMN_red, DUF4405 or Flavodoxin were collected for further analyses. InterPro accession numbers, taxonomic assignments, and amino acid sequences of flavinylated candidates presented in this study were included in supplementary table 1.

### Protein model prediction by AlphaFold2

Predicted 3D models for selected flavinylated protein monomers or complexes were generated using AlphaFold2 and AlphaFold2-multimer (ColabFold v1.5.2) (Jumper et al., 2021; Evans et al., 2021; Mirdita et al., 2022). The resulting PDB files containing predicted structures were visualized, examined, or aligned in PyMol.

### In vitro confirmation of covalent FMN binding in flavinylated flavodoxins and FMN reductases

#### E. coli expression strains

DNA fragments containing wild-type or point mutant ORFs of flavodoxins (*Anaerocolumna xylanovorans*, NCBI accession SHO45324.1; *Desulfosporosinus lacus*, NCBI accession WP_073032509.1) and FMN reductases (*Lactococcus lactis*, NCBI accession WP_021723379.1; *Lactococcus taiwanensis*, NCBI accession WP_205872264.1) were synthesized using IDT gBlocks (Integrated DNA Technologies). Primers with overhanging sequences homologous to either 5’ or 3’ end of target gene fragments were used to linearize pMCSG53 expression vectors at the multiple cloning sites through PCR reactions (Q5 High-Fidelity 2X Master Mix, New England Biolabs). Amplicons were subsequently gel-extracted (Wizard SV Gel and PCR Clean-Up System, Promega), quantified, and combined with corresponding gene inserts in Gibson reactions (NEBuilder HiFi DNA Assembly Master Mix, New England Biolabs) to allow integration of targeted genes. Expression constructs were then transformed into *E. coli* BL21 and successful transformants were selected on LB agar containing 100 mg/mL of carbenicillin. LB cultures of transformant colonies were supplemented with 15% w/v glycerol and stored in -80°C until use.

#### Purification of FMN transferase ApbE from Listeria monocytogenes

To ensure consistent flavinylation activity, we developed an *in vitro* flavinylation assay using the previously characterized FMN transferase ApbE protein encoded by *Listeria monocytogenes* 10403S, thereafter referred to as lm_ApbE. *E. coli* BL21 expression strains containing pMCSG53::*lm_apbe* expression constructs were generated through steps similar to those mentioned above with 6xHis-tag at the N-terminus. To purify lm_ApbE, overnight cultures of the expression strain were diluted to an optical density of OD600 = 0.05 in 1 L of LB and incubated at 37°C with aeration. After 2 h of incubation, a final concentration of 1 mM of IPTG was added to allow induction of lm_ApbE expression at 30°C overnight. Cell pellet was then collected through centrifugation at 7,000 x g for 15 min and frozen at -80°C overnight. Cell pellet was resuspended in a solution containing 50mM of Tris-HCl pH = 7.5, 300 mM of NaCl, and 10 mM of imidazole at a volume that is 5 times the weight of cell pellet. Resulting mixture was lysed through sonication (8 x 30 s pulses) and cleared by centrifugation 40,000 x g for 30 min. Supernatants of cell lysates were passed through a nickel bead column (Profinity IMAC Ni-Charged Resin, Bio-Rad) to allow binding of lm_ApbE-6xHis, which was then eluted with 500 mM imidazole. Successful elution of lm_ApbE-6xHis were confirmed through 12% SDS-PAGE. Filtrate samples were then purified using a ÄKTA pure chromatography FPLC system (Cytiva), and elution fractions containing lm_ApbE-6xHis were subsequently concentrated (4,000 x g; Pierce Protein Concentrators PES, 10K MWCO, Thermo Scientific) and quantified using a spectrophotometer (DS-11 FX+ spectrophotometer, DeNovix).

#### In vitro expression and flavinylation of flavodoxin and FMN reductase candidates

To confirm in vitro covalent binding of FMN on target candidate proteins, overnight cultures of *E. coli* BL21 strains containing corresponding expression vectors mentioned above were re-inoculated in 3 mL of LB and grown in presence of 1 mM IPTG with aeration at 30°C overnight. Overnight cultures were then diluted to an optical density of OD600 = 0.5 and centrifuged for 1 min at 21,100 x g. Resulting cell pellets were resuspended in 100 uL of lysis buffer (500 ug/mL of lysozyme, 300 mM of NaCl, and 10 mM of imidazole in 50mM of Tris-HCl pH = 7.5) and incubated on ice for 30 min. Cell lysates were then combined with 0.3 uM of lm_ApbE, 1 mM of FAD, and 5 mM of MgSO_4_ and incubated at 4°C overnight with rotation to enable flavinylation. Reaction mixtures were then separated into aqueous or solid phases by centrifugation at 21,100 x g for 1 min and subsequently incubated at 98°C for 10 min. Both aqueous and solid portions (resuspended in 100 uL of lysis buffer) were then run on 12% SDS-PAGE. To confirm successful flavinylation, we leveraged the UV resonance property of the isoalloxazine ring of the FMN moiety, which led to a bright band at expected molecular weight for targeted proteins when the SDS-PAGE gel is visualized under UV (iBright 1500 imaging system, Invitrogen).

#### In vitro confirmation of heme-binding activity in FezC and DUF4405

Cloning, expression, and purification of FezC (*Desulfurispirillum indicum*, WP_013506634.1) or DUF4405 (*Oscillospiraceae bacterium*, MBD5117352.1) were done in similar procedures as lm_ApbE, except that solid phase of cell lysates was used for downstream purification because FezC and DUF4405 are membrane proteins. Proteins in the solid phase of cell lysates were solubilized using a previously published protocol (Kupke et al., 2020). Briefly, pelleted cell lysates were resuspended in a solution containing 50mM of Tris-HCl pH = 7.5, 300 mM of NaCl, 10 mM of imidazole, and 1% w/v LDAO, and were subsequently purified as previously described using nickel bead column and FPLC (eluted with 500 mM imidazole + 0.1% w/v LDAO). Heme binding activity of purified FezC or DUF4405 was confirmed using a previously published protocol for pyridine hemochromagen assay (Barr et al., 2016). Briefly, samples containing 1 mg/mL of purified FezC or DUF4405 were mixed with a solution containing 0.2 M NaOH, 40% (v/v) pyridine, and 500 μM potassium ferricyanide to oxidize protein samples. Oxidized proteins were then measured for their absorbance at 300 - 700 nm. Samples were then combined with a reducing solution containing 0.5 M sodium dithionite in 0.5 M NaOH to acquire reduced FezC or DUF4405, which were then similarly examined for its absorbance at the same range of wavelength.

## ACKNOWLEDGMENTS

Research reported in this publication was supported by funding from the National Institutes of Health (K22AI144031 & R35GM146969 to S.H.L) and the Searle Scholars Program (to S.H.L).

